# Zhx2 is a candidate gene underlying oxymorphone metabolite brain concentration associated with state-dependent oxycodone reward

**DOI:** 10.1101/2022.03.18.484877

**Authors:** Jacob A. Beierle, Emily J. Yao, Stanley I. Goldstein, William B. Lynch, Julia L. Scotellaro, Katherine D. Sena, Alyssa L. Wong, Colton A Linnertz, Olga Averin, David E. Moody, Christopher A. Reilly, Gary Peltz, Andrew Emili, Martin T. Ferris, Camron D. Bryant

## Abstract

Understanding the pharmacogenomics of opioid metabolism and behavior is vital to therapeutic success as mutations can dramatically alter therapeutic efficacy and addiction liability. We found robust, sex-dependent BALB/c substrain differences in oxycodone behaviors and whole brain concentration of oxycodone metabolites. BALB/cJ females showed robust state-dependent oxycodone reward learning as measured via conditioned place preference when compared to the closely related BALB/cByJ substrain. Accordingly, BALB/cJ females also showed a robust increase in brain concentration of the inactive metabolite noroxycodone and the active metabolite oxymorphone compared to BALB/cByJ mice. Oxymorphone is a highly potent full agonist at the mu opioid receptor that could enhance drug-induced interoception and state-dependent oxycodone reward learning. Quantitative trait locus (**QTL**) mapping in a BALB/c F2 reduced complexity cross revealed one major QTL on chromosome 15 underlying brain oxymorphone concentration that explained 32% of the female variance. BALB/cJ and BALB/cByJ differ by fewer than 10,000 variants which can greatly facilitate candidate gene/variant identification. Hippocampal and striatal cis-expression QTL (eQTL) and exon-level eQTL analysis identified *Zhx2*, a candidate gene coding for a transcriptional repressor with a private BALB/cJ retroviral insertion that reduces Zhx2 expression and sex-dependent dysregulation of CYP enzymes. Whole brain proteomics corroborated the Zhx2 eQTL and identified upregulated CYP2D11 that could increase brain oxymorphone in BALB/cJ females. To summarize, Zhx2 is a highly promising candidate gene underlying brain oxycodone metabolite levels. Future studies will validate *Zhx2* and its site of action using reciprocal gene editing and tissue-specific viral manipulations in BALB/c substrains.

**Significance Statement:** Our findings show genetic variation can result in sex-specific alterations in whole brain concentrations of bioactive opioid metabolites following oxycodone administration, and reinforces the need for sex as a biological factor in pharmacogenomic studies. The co-occurrence of female-specific increased oxymorphone and state-dependent reward learning suggests that this minor yet potent and efficacious metabolite of oxycodone could increase opioid interoception and drug-cue associative learning of opioid reward which has implications for cue-induced relapse of drug-seeking behavior.

## INTRODUCTION

Opioids are analgesics prescribed for severe pain and have a high addiction liability. The heritability of opioid use disorder is estimated to be between ∼25-70% (Chan et al., 2011; Goldman et al., 2005; Kendler et al., 2000, 2003; Tsuang et al., 1996, 1998). Genome wide association studies have identified a handful of candidate genes contributing to the heritability of OUD in humans (Crist et al., 2019) but we still know very little regarding the genetic contributions to opioid use disorder.

Genetic variation can influence the pharmacokinetic properties of xenobiotics, such as opioids, and in turn, the therapeutic potency, efficacy, and addiction liability. Oxycodone is a commonly prescribed semisythetic opioid that is metabolized via phase I metabolism into biologically inactive noroxycodone (∼45% via CYP3A4) and the biologically active oxymorphone (∼19% via CYP2D6) (Huddart et al., 2018). Oxymorphone exhibits greater potency and efficacy at activating the mu opioid receptor than Oxycodone (Lalovic et al., 2006; Thompson et al., 2004). An unresolved question is whether oxymorphone contributes to individual differences in the analgesic and addictive properties of oxycodone (Andreassen et al., 2012; Candiotti et al., 2009; Lemberg et al., 2010; McMillan et al., 2019; Zwisler et al., 2009, 2010). Given the widespread use and misuse of oxycodone, the contribution of oxymorphone to oxycodone’s motivational and therapeutic properties is an important question.

The use of mice in unbiased, discovery-based genetic analysis of opioid traits provides complimentary advantages to human genetic studies by allowing control of genetic background, precise dosing, collection of end point tissues at appropriate time points, and the ability to validate causal variants through gene editing. Inbred mouse strains show large variation in opioid behaviors (Belknap et al., 1993, 1995; Bergeson et al., 2001; Bubier et al., 2020; Kest et al., 2002; Solecki et al., 2009; Wilson et al., 2003), and forward genetic studies have identified candidate genes influencing such traits as opioid-induced respiratory depression (*Galnt11*: Bubier et al., 2020), locomotor stimulation (*Csnk1e*: Bryant et al., 2009, 2012), antinociception (*Kcnj9*: Smith et al., 2008; Car8: Levitt et al., 2017), physiological withdrawal (*Gnao1*: Kest et al., 2009); *CNIH3*: Nelson et al., 2016), tolerance, and hyperalgesia (*Mpdz*: Donaldson et al., 2016).

More recently, the use of near-isogenic substrains, distinguished primarily by genetic variants that have arisen from genetic drift, have proven to be quite useful for genetic mapping of complex traits in “Reduced Complexity Crosses” (Bryant et al., 2018; 2020). Large phenotypic variance combined with a drastic reduction in the density of genetic polymorphisms facilitates the identification of causal genes/variants that can readily be validated via gene editing (Mulligan et al., 2019). Reduced complexity crosses have been used between multiple mouse substrains (e.g., C57BL/6, DBA/2, BALB/c) to map the genetic basis of complex phenotypes such as psychostimulant sensitivity, binge-like eating, and thermal nociception (Beierle et al., 2022; Goldberg et al., 2021; Harkness et al., 2015; Kirkpatrick et al., 2017; Kumar et al., 2013; Miner et al., 2017; Reed et al., 2018; Shi et al., 2016).

Regarding this study, BALB/cJ **(J)** and BALB/cByJ **(By)** substrains were separated in 1935 after the F37 generation of inbreeding and have since been maintained as separate substrains. Fewer than 10,000 SNPs and indels are estimated to distinguish J from By, comprising a 500-fold reduction in genetic complexity compared to C57BL/6J versus most classical inbred strains (Keane et al., 2011; Yalcin et al., 2011). The J and By substrains differ in several interesting neurobehavioral phenotypes (Dam et al., 2019; Hilakivi & Lister, 1989; Jager et al., 2020; Perincheri et al., 2005; Poyntz et al., 2019; Sittig et al., 2014, 2014; Turner et al., 2008; Velez et al., 2010). Given this phenotypic variation, BALB/c substrains likely harbor readily identifiable causal genetic variants mediating important behavioral traits and we have previously employed a reduced complexity cross in BALB/c substrains to map QTLs, eQTLs, and candidate genes underlying thermal nociception and brain weight (Beierle et al., 2022).

In this study we observed robust sex-dependent BALB/c substrain differences in state-dependent expression of oxycodone reward learning in a conditioned place preference paradigm. We also observed robust, sex-dependent strain differences in brain concentrations of oxycodone (**OXY**) and its metabolites noroxycodone (**NOR**) and oxymorphone (**OMOR**). To identify candidate genes underlying brain OXY and metabolite concentration, we conducted metabolite quantitative trait locus (**QTL**) mapping and gene-level and exon-level expression QTL (**eQTL**) mapping in a BALB/c reduced complexity cross as well as complementary whole brain proteomic and liver transcriptomic analysis. The combined results strongly implicate *Zhx2* as a candidate gene underlying brain concentration of OMOR.

## METHODS

### Mice

All experiments were conducted in accordance with the National Institutes of Health Guidelines for the Use of Laboratory Animals (8^th^ Ed.) (National Research Council (US) Committee for the Update of the Guide for the Care and Use of Laboratory Animals, 2011) and were approved by the Institutional Animal Care and Use Committee at Boston University School of Medicine. BALB/cJ (**J**) and BALB/cByJ (**By**) mice (7 weeks old) were purchased from The Jackson Laboratory (Bar Harbor, ME; #000651, #001026), housed four per cage, and allowed six days to acclimate before testing. BALB/cJ and BALB/cByJ mice were also bred in house for select parental strain experiments and were always conducted with mice ordered from Jackson Laboratory to avoid the fixation of genetic drift in our breeding population. These mice were tested between the ages of 47 and 83 days old. BALBcJ x BALBcByJ-F1 and -F2 mice were bred in house as described below and were tested between the ages of 56 and 134 days old.

The age range was larger than our typical range of 50-100 days old because of the COVID-19 pandemic. All mice were maintained on Teklad 18% protein diet (Envigo, Indiana; #2018) and were tested from 1030-1300 in the light phase of a 12 h light/dark cycle (lights on at 0630).

### Drugs

Oxycodone hydrochloride was purchased from Sigma-Aldrich (St. Louis, MO USA), was dissolved in sterilized saline (0.9% NaCl), and was administered in an injection volume of 10 ul/g.

### Conditioned place preference in BALB/c substrains

BALB/cJ and BALB/cByJ substrains for parental strain experiments were trained and tested for conditioned place preference using a nine-day protocol in a two-chamber apparatus (20 cm x 40 cm) as previously described (Kirkpatrick & Bryant, 2015). On Day 1, mice were administered saline (**SAL**), placed into the left side, and provided open access to both sides to assess initial preference. On Days 2 and 4 mice were confined to the right chamber and administered 1.25 mg/kg OXY (i.p.) or volume-matched SAL (10 ul/g, i.p.). The left and right sides were distinguished by different plastic floor texture inserts. On Day 3 and Day 5, mice were confined to the left chamber and administered SAL (i.p.). Drug-free preference was measured on Day 8 following a SAL injection (i.p.), placement into the left side, and providing open access to both sides for 30 min. State-dependent preference was measured on Day 9 which was procedurally identical to Day 8 except that (OXY (1.25 mg/kg, i.p.) was injected to OXY-trained mice and SAL (i.p.) to SAL-trained mice. Preference was defined as the difference in time spent on the drug-paired side between Day 1 and either Day 8 or Day 9. All testing started at 1100, all training and testing sessions lasted 30 min, and all testing apparatuses were housed in unlit sound attenuating chambers. Video was captured with overhead cameras (Swann Security Inc, Santa Fe Springs, CA, USA) and graded using ANYMAZE tracking software (Wood Dale, IL, USA). These experiments were conducted in 9 cohorts, and with separate distinct aims.

To establish behavioral differences between substrains, 3 cohorts totaling 88 mice were ordered from The Jackson Laboratory was used to assess several OXY related phenotypes over 7 weeks. The first 9 days of this testing was CPP as described above, and for brevity and specificity we have omitted subsequent testing. Three subsequent cohorts of inbred BALB/cJ and BALB/cByJ substrains were bred in house using mice ordered from The Jackson Laboratory to ensure observed differences in behaviors could be replicated in mice bred at Boston University, as all F2 mice would be generated in our facilities. This group constitutes 77 mice from 47 – 83 days old at day one of testing. Finally, 3 more cohorts totaling 120 mice, ordered at 7 weeks old from The Jackson Laboratory, were tested for CPP and subsequently tissues were taken for scRNA-seq (data not included), whole brain proteomics, and liver RNA-seq and proteomic testing.

### Samples generated for assessment of whole brain OXY, NOR, and OMOR concentrations

Whole brain concentrations were assayed in BALB/cJ and BALB/cByJ substrains (20 J, 20 By) from two different cohorts with slightly different protocols. The first cohort consisted of 16 mice [7 J (4 females, 3 males), 9 By (6 females, 3 males)] bred in house and 56-82 days old, that were tested using a three-day protocol. On Day 1 and Day 2, mice were administered saline (10 ul/g, i.p.) and placed in a 20 cm x 40 cm open arena (the CPP chambers minus the floor textures and divider) to allow for habituation to injection and context. On Day 3, mice were administered 1.25 mg/kg OXY (i.p.) and recorded for 30 min within the apparatus. Immediately following testing, mice were sacrificed by rapid decapitation, brains were dissected from the skull, olfactory bulbs trimmed, and brainstem trimmed at the pons-medulla boundary. Brains were then flash frozen in a bath of alcohol and dry ice, weighed, and stored at -80°C until processed. All testing started at 11:00, all testing sessions lasted 30 min, and all testing apparatuses were housed in sound attenuating chambers. Video was captured with overhead cameras and graded using ANYMAZE tracking software.

The second cohort of mice comprised 25 OXY-treated mice [13 J (6 females, 7 males), 12 ByJ (6 females, 6 males)] that were trained for state-dependent OXY-CPP (the same samples were used for the liver RNA-seq experiment that is described below). Immediately after Day 9 of CPP (30 min after the third dose of OXY; 1.25 mg/kg, i.p.), brains were dissected as described above and stored at -80°C before analysis. Thus, the two cohorts used for metabolite analysis differed with regard to the number of OXY injections (one versus three) and the size of the arena in which they were placed (20 cm x 40 cm or 20 cm x 20 cm).

### Oxycodone and metabolite sample preparation

Frozen and pre-weighed mouse brains were shipped from Boston University School of Medicine to the University of Utah Center for Human Toxicology on dry ice. The samples were stored at -30°C until processed. Briefly, the samples were removed from the freezer in groups of 10 and and kept on ice. The brain sample was then transferred to a 15-mL polypropylene centrifuge tubeand 4 mL of Type 1 water was added to the tubeThe samples were then homogenized using the Sonics Vibra Cell sonicator fitted with the microtip at 30% power. Homogenization was done in three 15-s cycles with approximately 2 s between the cycles, resulting in ca. 4.5 mL of brain homogenate. After all 10 samples were sonicated, 1 mL of each homogenate was aliquoted into an individually marked 16x100 silanized glass tube. After these operations were completed, all test tubes were placed back in a freezer at -30 °C until further processing and analysis.

### Quantification of Oxycodone and metabolites by liquid chromatograohy-mass spectrometry

OXY, NOR, and OMOR concentrations were measured using a validated liquid chromatographytandem mass spectrometry (LC/MS/MS) method (Fang et al., 2013). Deuterated OXY-d6, NOR-d3, and OMOR-d3 were added as internal standards to each homogenate sample and mixed. The samples were then extracted under basic conditions (100 µL of concentrated ammonium hydroxide) with 4 mL of freshly prepared n-butyl chloride:acetonitrile (4:1 v/v). Briefly, the samples were vortex mixed for ∼30 seconds and then centrifuged at 1200 x*g* for 10 min. The upper organic layer of each sample was then transferred into a clean silanized glass tube and evaporated to dryness under a stream of filtered air at 40°C. The extracts were reconstituted in 75 µL of 0.1% formic acid in water, centrifugedto clarify, and the supernatants transferred into autosampler vials. The LC/MS/MS system was an Agilent 1100 series HPLC coupled to a Thermo Scientific TSQ Quantum Access Triple Stage Quadruple mass spectrometer. A YMC-Pack ODS-AQ 5µm 2.0x100 mm column (Waters, Milford, MA) was used for analysis, and the mass spectrometer was run in positive electrospray mode. The concentrations of oxycodone (**OXY**), noroxycodone (**NOR**), and oxymorphone (**OMOR**) were determined from the ratio of the peak area of each drug to the peak area of its internal standard, and comparison with the calibration curve that was generated from the analysis of human plasma fortified with known concentrations of OXY, NOR, OMOR, and their internal standards. The lower limit of quantitation of the assay for all analytes was 0.2 ng/mL (dynamic range of 0.2 to 250 ng/mL).

### F2 breeding and genotyping

BALB/c F2 mice were generated as described (Beierle et al., 2022). Testing began between 56 and 134 days old, a range increased beyond our normal testing window of 50-100 days because of the COVID19 shutdown. Genotypes were determined using the miniMUGA microarray (Sigmon et al., 2020), and we refined these results to 304 which were polymorphic and reliable between our BALB/c substrains.

### Whole genome sequencing and genotype calling of BALB/c substrains

We used existing whole genome sequences for BALB/c substrains for variant identification as we previously reported (Beierle et al., 2022).

### F2 mice for metabolite phenotyping

F2 mice were phenotyped for CPP as described above. Three days after the completion of testing, mice were tested for baseline hot plate latencies as previously described (Beierle et al., 2022), and subsequently injected with OXY so that the samples could be assayed for metabolism. At 12:00, following hot plate testing, saline control (OXY naïve) F2 mice who had previously received three SAL injections during the CPP experiment were administered an acute OXY injection (1.25 mg/kg i.p.), placed in a clean mouse cage for 30 min, then brains were collected, and flash frozen as described above. Sample processing and analysis was conducted as previously described.

### QTL mapping of brain OXY and metabolite concentrations

QTL mapping in the F2 cross was conducted as reported (Beierle et al., 2022). Briefly, poor quality markers and animals were excluded based on missing calls, non-mendelian inheritance, and inappropriate cross over counts within a mouse. After QC there were 133 F2 mice (68 F, 65 males) and 218 polymorphic markers from the miniMUGA with which to conduct QTL mapping for whole brain oxycodone and metabolites. To determine the QTL model, a three-way ANOVA considering Sex, Cohort, and Age on brain [OMOR] revealed a significant effect of Sex, but no other main effects or interactions. We therefore included Sex as an additive covariate in the model. Calculating QTLs using Haley-Knott regression was conducted as described (Beierle et al., 2022). Given the sparsity of miniMUGA markers within our chromosome 15 QTL interval, we designed an additional marker using the TaqMan fluorescent genotyping assay (Thermofisher Scientific). We targeted rs264203947 (chr15:57104609 bp). The assay was conducted using a 10 uL reaction consisting of 5 ng of DNA and 0.5 uL of the TaqMan assay in 1x TaqMan Genotyping Master Mix (ThermoFisher Scientific, 4371355) and run using the following cycle in a 12K QUANT STUDIO 12K FLEX (Thermofisher Scientific): 1x 95°C for 10 min, 40x 95°C for 15 s, 60C 1 min. Power analysis of F2 whole brain [OMOR] was conducted using the package R/QTLdesign (Sen et al., 2007). Power analysis illustrating effect sizes versus sample sizes required were computed for Day 9 preference in R/QTLdesign (Sen et al., 2007) using the effect size derived from the parental strain differences.

### Gene-level, exon-level expression QTL (eQTL) mapping in F2 mice

A subset of 64 F2 mice were sacrificed for eQTL analysis after D9 CPP testing as described in Beierle et al., 2022. Briefly, we analyzed hippocampal and striatal RNA-seq counts using R/MatrixEQTL (Shabalin, 2012) using the ‘linear cross’ model and included Sex, RNA extraction Batch (RNA extraction), and Prior Treatment (SAL, OXY) as covariates. The same model was used to analyze intron and exon feature counts from the R/ASpli (Mancini et al., 2021), as described (Beierle et al., 2022).

### Whole brain peptide proteomics in BALB/c parental substrains

24 BALB/cJ (**J**) and 24 BALB/cByJ (**By**) mice (12 per sex per substrain) were trained and tested for drug-free and state-dependent CPP using 1.25 mg/kg OXY (i.p.) and published procedures (Kirkpatrick & Bryant, 2015) that are summarized above. Immediately after testing for state-dependent CPP on Day 9 (30 min post-OXY), mice were sacrificed and 16 brains were harvested for proteomic analysis as described in Beierle et al., 2022. Briefly, 8 BALB/cJ (2/sex/tx) and 8 BALB/cByJ (2/sex/tx) whole brains were collected, homogenized, fractionated, and MS/MS spectra acquired. Spectra were searched against the complete SwissProt mouse proteome, MaxQuant was used for data normalization, and in-house scripts were used for statistical analysis. A complete list of differentially expressed proteins can be found in Beierle et al., 2022.

### Parental strain Liver dissection, extraction, RNAseq, and DEG analysis

16 BALB/cJ (8 females, 8 males) and 16 BALB/cByJ mice (8 females, 8 males) were trained and tested for state-dependent OXY-CPP using 1.25 mg/kg OXY (i.p.) as described above and sacrificed immediately after day 9 testing (30 min post-OXY) by rapid decapitation. The left lobe of the liver was harvested and submerged in RNA Stabilization Solution RNA*later^TM^* Solution (Thermofisher) and stored at 4°C. Briefly, RNA was extracted using Trizol and spin columns, sequenced on an Illumina NovaSEQ6000, then fastq files demultiplexed, trimmed, and aligned to the mm10 mouse reference genome (Ensembl). We assessed the effect of Substrain on differential gene expression in the Sex-collapsed dataset, while controlling for the effects of Prior Treatment. We also conducted separate analyses in females and males while controlling for the effect of Treatment. Full details describing RNA extraction and data processing can be found in Beierle et al., 2022.

### Liver proteomics in BALB/c parental substrains

The same 16 SAL treated mice used for liver RNA seq were also used for liver proteomics (8 BALB/cJ, 4M 4F; 8 BALB/cByJ mice 4M, 4F). The left lateral lobes of the liver were homogenized, fractionated, and MS/MS spectra acquired. Livers were processed and analyzed in the same manner as the whole brain samples above Notable, one mouse (female By) was excluded from analysis because the tandem mass tag multiplexing channel corresponding to this sample was empty.

## Results

### BALB/cJ mice show increased state-dependent OXY-CPP compared to BALB/cByJ mice

The CPP protocol and apparatus are pictured in **Figure 1A**. In examining drug-free OXY-CPP on Day 8 using a 3 way ANOVA, there was a significant main effect of Treatment (F(1,276)=5.8, **#**p = 0.019,), indicating a significant preference for OXY. However, there was no effect of Substrain (F(1,276)=1.9, p = 0.17), Sex (F(1,276)=1.92, p = 0.16), or any interactions (p = 0.12 – 0.74; **Supplementary Figure 1**). One video recording of the 285 recordings was lost for Day 8 and was therefore not in the present analysis.

**Figure 1:**
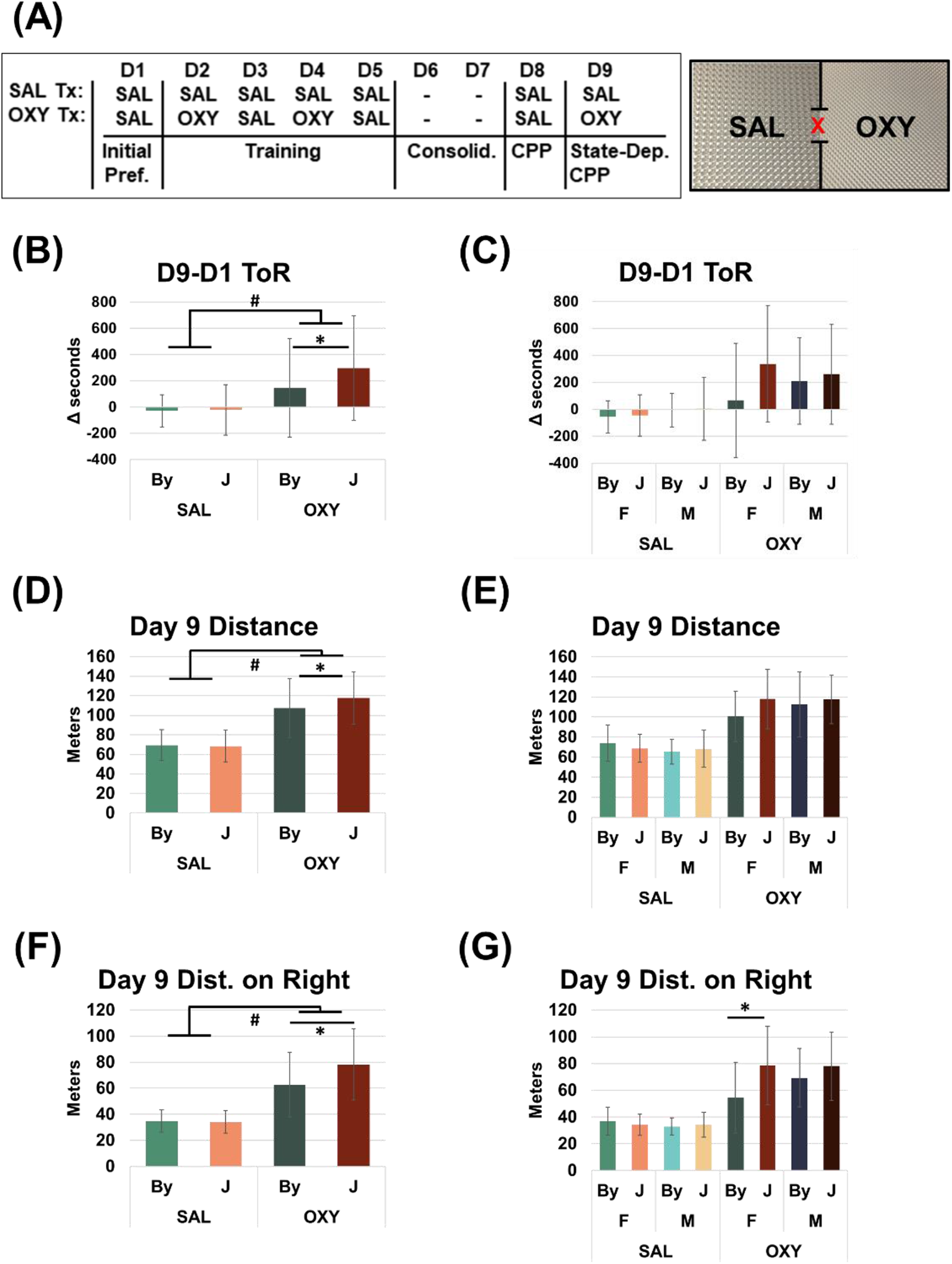
BALB/cJ (J) mice show enhanced state-dependent learning of OXY-CPP (1.25 mg/kg, i.p.) and concomitant locomotion compared to BALB/cByJ (By) mice. (**A**): Schematic of the CPP regimen and apparatus. (**B**): Results were analyzed using a three way ANOVA considering Substrain, Treatment, and Sex, error bars represent standard deviation. There was a Treatment x Substrain interaction (p = 0.048) that was explained by J mice showing greater OXY-CPP than By mice (Tukey post-hoc *p = 0.015). (**C**): Female mice accounted for a majority of the substrain difference in OXY-CPP. (**D**): For total OXY-induced locomotor activity, there was a significant Treatment x Substrain interaction (p = 0.043) that was explained by J mice showing greater OXY-induced locomotion than By mice (Tukey’s post-hoc *p = 0.037). (**E**): Females accounted for a majority of the substrain difference in total OXY-induced locomotor activity. (**F**): For OXY-induced locomotor activity specifically on the OXY-paired side (right side), there was a significant Treatment x Substrain interaction (p = 0.043) that was explained by J mice showing greater OXY-induced locomotor activity than By mice (Tukey’s post hoc *p = 0.037). (**G**): When Sex was included in the ANOVA model, there was a significant Treatment x Substrain x Sex interaction (p = 0.035) that was driven by increased OXY-induced locomotor activity in J females compared to By females (Tukey’s post hoc *p = 2.3e-5). Sample sizes were, from left to right in panel B: 32, 37, 36, 32, 29, 38, 37, and 43.

In examining state-dependent OXY-CPP on Day 9 using a 3 way ANOVA considering Substrain, Treatment, and Sex, there was a main effect of Treatment (F(1, 276)=51.06, **#**p=8.01e-12), indicating the presence of overall state-dependent CPP while under the influence of OXY (1.25 mg/kg, i.p.; **Figure 1B**). There was also a main effect of Substrain (F(1, 276)=5.13, p=0.024; **Figure 1B**), and an interaction between Treatment and Substrain (F(1, 276)=3.92, p=0.048). Tukey post-hoc test revealed a significant increase in state-dependent OXY-CPP in the OXY J versus the OXY By group (**\***adjP = 0.015). Adding Sex to the model did not reveal any significant main effect of Sex (F(1, 276)=1.03, p=0.31) or interactions between Sex and Strain (F(1, 276)=2.53, p=0.11), Sex and Treatment (F(1, 276)=0.17, p=0.68) or three way interaction (F(1, 276)=2.37, p=0.12). Nevertheless, to facilitate later comparison with the metabolite results, we also present the same data stratified by Sex (**Figure 1C**). One video recording of the 285 recordings was lost for Day 9 and is therefore not in the present analysis.

Analysis of OXY-induced locomotor activity (m) during state-dependent CPP assessment on Day 9 revealed a main effect of Treatment (F(1, 276)=258.52, **#**p=2e-16), a trending effect of strain (F(1, 276)=3.2, p=0.075) and an interaction between Treatment and Substrain (F(1, 276)=4.78, p=0.042). Tukey’s post-hoc test revealed a significant increase in J versus By OXY groups (**\***adjP = 0.037; **Figure 1D**). When considering Sex, we observed no main effect (F(1, 276)=0.05, p=0.82) nor a Treatment x Sex interaction (F(1, 276)= 3.08, p=0.08) or Treatment x Substrain x Sex interaction (F(1, 276)= 3.356, p=0.068; **Figure 1E**).

Analysis of Day 9 locomotion on the right side revealed a main effect of Treatment (F(1, 276)=251.04, **#**p=2e-16), a main effect of Substrain (F(1, 276)=10.75, p=0.001) and interaction between Treatment and Substrain (F(1, 276)= 10.78, p=6.3e-4; **Figure 1F**). Concerning Sex, we observed no main effects (F(1, 276)=1.12, p=0.29), but did observe a significant Treatment x Substrain x Sex interaction (F(1, 276)= 4.485, p=0.035). Tukey’s post hoc test revealed that the three-way interaction was driven by a significant increase in OXY-induced locomotor activity on the right side in OXY J females versus OXY By females (**\***adjP = 2.3e-5; **Figure 1G**).

### Increased whole brain [OXY], [NOR], and [OMOR] in BALB/c J versus BALB/cByJ mice

Given the differences in testing protocol between the two cohorts used for parental strain whole brain concentrations, we analyzed the data separately and we observed the same overall pattern of results. For cohort 1, analysis of whole brain [OXY] revealed a significant effect of Substrain (F(1, 12)= 8.3, p=0.014) and interaction between Substrain and Sex (F(1, 12)= 5.68, p=0.035, **Figure 2A**), but no effect of Sex (F(1, 12)= 0.16, p=0.7). Tukey’s post hoc test revealed this interaction to be mediated by a significant increase in whole brain [OXY] between the Males (**\***p = 0.015) not present in Females (p = 0.85, **Figure 2A**). For whole brain [NOR], there was a significant effect of Substrain (F(1, 12)= 7.88, p=0.016), Sex (F(1, 12)= 14.72, p=0.002), and a trending interaction (F(1, 12)= 3.3, p=0.094, **Figure 2B**). Tukey’s post hoc test revealed a significant increase in whole brain [NOR] in the J Females compared to all other groups (**\***p = 0.016 – 0.003, **Figure 2B**).

**Figure 2:**
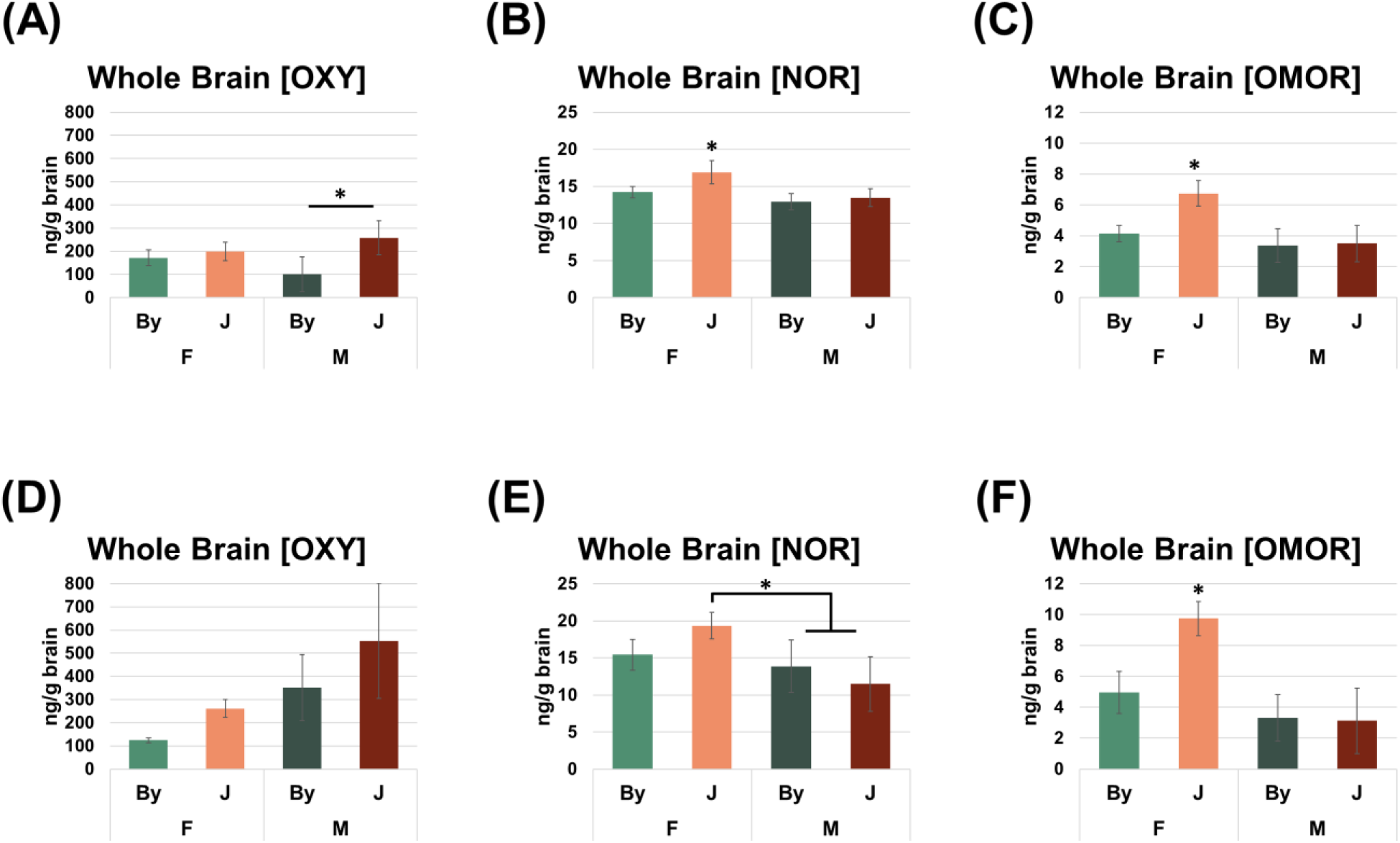
BALB/cJ mice show elevated whole brain concentrations of OXY, NOR, and OMOR compared to BALB/cByJ mice at 30 min post-injection of OXY (1.25 mg/kg, i.p.) (**A-C**): All analysis was conducted using a 2 way ANOVA considering Substrain and Sex, error bars represent standard deviation. For the first cohort of mice, we observed an effect of Substrain on brain concentrations of all compounds. We also detected significant main effects of Sex and interactions of Sex with Substrain for [NOR] (p = 0.048) and [OMOR] (p = 0.005). In both cases this effect was driven by increased drug concentrations in J females (Tukey post-hoc *p = 0.022 – 9.9e-4). Sample sizes were, from left to right in panel A: 6, 4, 3, and 3. (**D**): For the second cohort of mice in which there were slightly different procedures (see Methods), for whole brain [OXY], we observed an effect of Substrain (p = 0.007) and Sex (p = 3.1e-4), but no interaction. (**E**) For whole brain [NOR], there was no effect of Substrain, but there was a significant effect of Sex (p = 5.2e-4) and a Sex x Substrain interaction (p = 0.014) that was driven by a significant increase in J females vs. J males (Tukey’s post-hoc *p = 5.2e-4). (**F**) Finally, for whole brain [OMOR], there were main effects of Substrain (p = 0.004) and Sex (p = 1.5e-6) and a Substrain x Sex interaction (p = 8.7e-4) that was driven by the increase in [OMOR] in J females compared to all three other groups (Tukey’s post-hoc *p < 2.1e-4). Sample sizes were, from left to right in panel A: 6, 6, 6, and 7.

Analysis of whole brain [OMOR] revealed a main effect of Substrain (F(1, 12)= 11.48, p=0.005), Sex (F(1, 12)= 19.18, p=7e-4), and an interaction (F(1, 12)= 7.86, p=0.016,) that was also driven by an increase in brain [OMOR] in J females compared to the three other groups (*p = 0.002 – 0.001; **Figure 2C**).

For whole brain [OXY] in cohort 2, we observed a main effect of Substrain (F(1, 21)= 8.9, p=0.007) and Sex (F(1, 21)= 18.6, p=3.1e-4), but no interaction (F(1, 21)= 0.28, p=0.6, **Figure 2D**). Analysis of whole brain [NOR] revealed a main effect of Sex (F(1, 21)= 16.8, p=5.2e-4), and a Substrain x Sex interaction (F(1, 21)= 7.13, p=0.014), but no main effect of Substrain (F(1, 21)= 0.151, p=0.7). Tukey post hoc test revealed that the Substrain x Sex interaction was driven by an increase in [NOR] in J females compared to J males (**\***p = 0.02; **Figure 2E**) and By males (*p = 5.1e-4), but not By females (p = 0.13). Finally analysis of whole brain [OMOR] revealed a significant effect of Substrain (F(1, 21)= 10.24, p=0.01), Sex (F(1, 21)= 43.7, p=1.5e-6), and a Substrain x Sex interaction (F(1, 21)= 15.04, p=8.7e-4) that was driven by an increase in J females compared to the three other groups (*p=2e-4 – 1.4e-5; **Figure 2F**).

### QTL model selection for whole brain [OMOR] in F2 mice

A genetic map of the 219 markers that passed QC measures is shown in **Supplementary Figure 2A**. Details concerning the TaqMan fluorescent genotyping assay are provided in **Supplementary Table 1.** To derive an appropriate model for QTL mapping, we considered the effects of Sex, Cohort, and Age on whole brain [OMOR] via ANOVA in our F2 mice and observed a main effect of Sex (F(1, 125)= 17.49, p=5.4e-5), but no effect of Cohort (F(1, 125)= 0.62, p=0.43), Age (F(1, 125)= 0.064, p=0.8), or any interactions (ps > 0.49; **Supplementary Figure 2B**). Therefore, we included Sex as an additive covariate in our QTL analysis.

### A major QTL on chromosome 15 underlying whole brain [OMOR]

We did not identify any genome-wide significant QTLs underlying BALB/c substrain differences in state-dependent OXY-CPP when considering treatment as an interactive covariate and sex as an additive covariate (**Supplementary Figure 3A,B**). This null result supports previous observations from our group and others that CPP is not a highly heritable trait (Bryant et al., 2014; Cunningham et al., 1991; Gonzales et al., 2018; Kirkpatrick et al., 2017; Philip et al., 2010; Ruan et al., 2020). Nevertheless, CPP is clearly a useful phenotype as it led us to measure a more heritable trait - brain concentration of OXY and its metabolites and to ultimately identify a plausible candidate gene that could influence both traits.

We identified a genome-wide significant QTL on chromosome 15 (LOD = 7.07; p<0.001) for whole brain [OMOR] that peaked at 30 cM (67 Mb) and explained 19% of the variance in all mice and 32% in females (**Figure 3A,B**, **Table 1**). Notably, we did not observe significant QTLs for whole brain [OXY] or [NOR]. At the peak associated marker for whole brain [OMOR] (rs264203947), the effect of Genotype and Sex recapitulated the parental substrain difference, in which females homozygous for the J allele showed an increase in whole brain [OMOR] (**Figure 3C**). **Table 1** summarizes the QTL mapping results. There are 55 polymorphic protein coding genes within the Bayes confidence interval (**Supplementary Table 2**).

**Figure 3:**
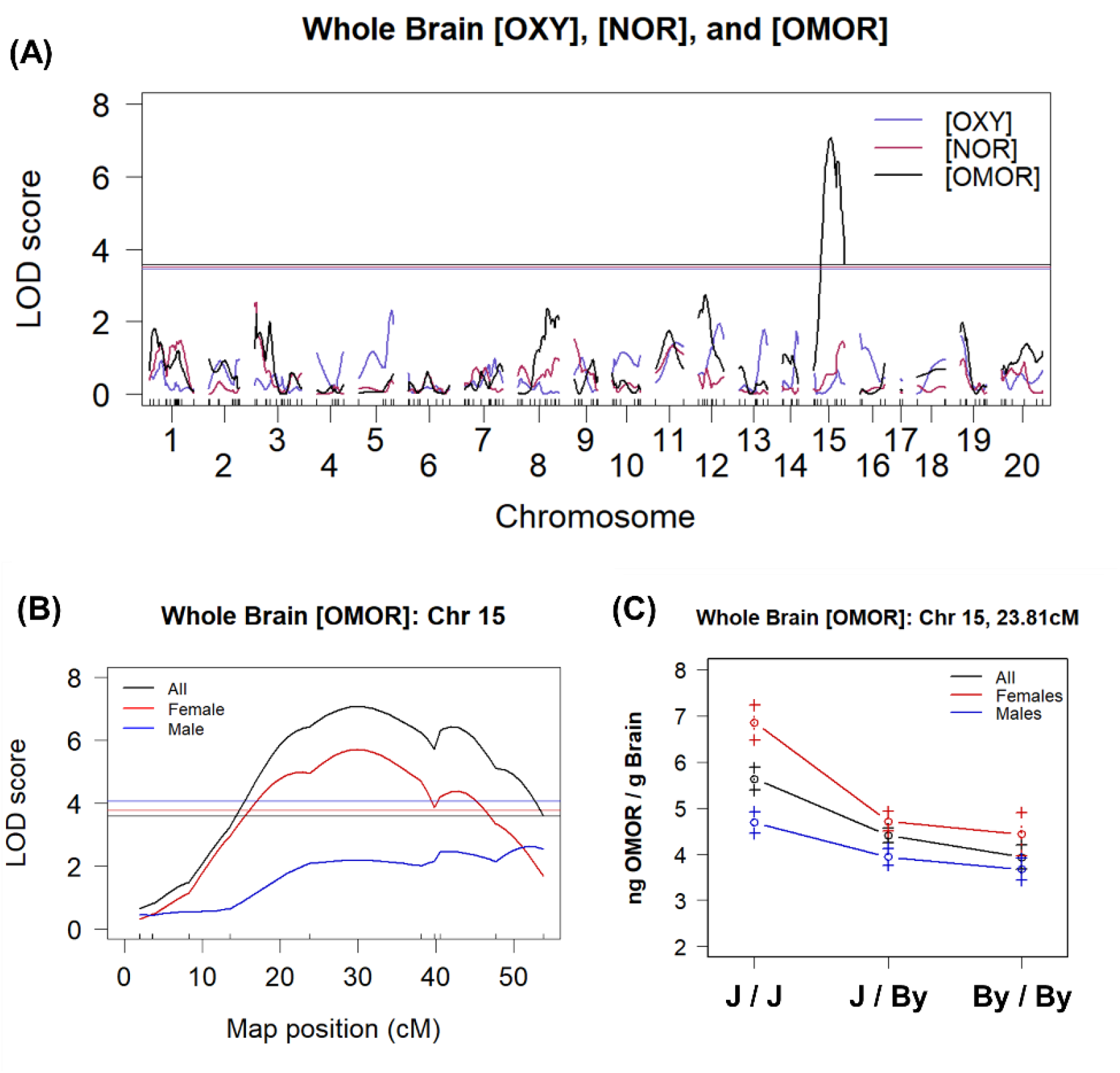
A major QTL on chromosome 15 underlies variation in whole brain OMOR concentration following OXY administration (1.25 mg/kg, i.p.) (**A**): In 133 F2 mice and considering Sex and additive covariate, a single genome-wide significant QTL peaked on medial chromosome 15 (LOD=7.07, p < 0.001) that explained 19% of the sex-combined phenotypic variance. (**B**): Chromosome 15 QTL plot shows a peak association at 30 cM (68 Mb). (**C**): At the peak-associated marker, F2 mice sorted by BALB/c genotype precisely recapitulated the result from the parental substrains (J > By) of increased whole brain [OMOR] in females with the homozygous J/J genotype versus the homozygous By/By genotype.

**Table 1:** Summary of the chromosome 15 QTL for whole brain [OMOR]. Summary statistics are provided for the sex-combined and females-only analysis.

| N | Mice | Phenotype | Model | Chr | Peak (cM) | Peak (Mb) | LOD | pVal | Bayesian<br>Interval<br>(cM) | Bayesian<br>Interval<br>(Mb) | 1.5 LOD<br>Drop<br>(cM) | 1.5 LOD<br>Drop (Mb) | % Var |
| --- | --- | --- | --- | --- | --- | --- | --- | --- | --- | --- | --- | --- | --- |
| 133 | All | Brain [OMOR] | Sex Add | 15 | 29.95 | 68.64 | 7.07 | <0.001 | 20.95 -<br>43.95 | 53.79 -<br>88.49 | 18.95 -<br>46.95 | 50.39 -<br>95.94 | 19.10 |
| 68 | Female | Brain [OMOR] | No Cov | 15 | 29.95 | 68.64 | 5.71 | <0.001 | 16.95 -<br>43.95 | 44.63 -<br>88.49 | 12.95 -<br>47.8 | 31.48 -<br>93.65 | 32.05 |

Power analysis using QTLdesign (Sen et al., 2007) indicated that using whole brain [OMOR] from 133 F2 mice, we had 80% power to detect additively inherited QTLs that explained 14% of phenotypic variance (**Supplementary Figure 4A**). Analysis of D9 preference revealed we were underpowered to map our observed P0 differences in preference, needing to test 311 OXY treated F2 mice compared to our 200 (**Supplementary Figure 4B**). Our current sample size could only successfully map QTLs explaining ∼12% of the observed variance (**Supplementary Figure 4C).**

### Transcript-level eQTL and exon-level eeQTL analysis of chromosome 15 identifies *Zhx2* as a candidate gene underlying whole brain [OMOR]

We obtained an average of 33.9 million paired-end reads across all 128 RNAseq samples from the 64 F2 mice (16/sex/treatment). **Table 3** shows a list of transcripts with significant eQTLs on chromosome 15 within the striatum and hippocampus (FDR < 0.05). In considering both striatal and hippocampal tissue, Zhx2 was the only transcript containing an eQTL (FDR < 0.05) that peaked with the same marker (24 cM) that was identified for whole brain [OMOR]. (Striatum FDR = 1.4e-20, Hippocampus FDR = 7.2e-8). A complete list of eQTL genes with unadjusted P values can be found in Beierle et al., 2022. QTL analysis at the exon/intron level identified 39 differentially used, non-binned, features within the striatum and 55 in the hippocampus associated with the peak marker for whole brain [OMOR] (FDR < 0.05). In both tissues Zhx2 features comprised the most significant cis-exon-level eQTL, and in both tissues introns 1 and 3 and exons 3 and 4 of Zhx2 were differentially regulated as a function of Genotype (**Table 3**). Interestingly, while the use of short read DNA sequencing failed to identify any variants assigned to Zhx2, there is a well-documented structural variant within intron 1 of this gene that decreases Zhx2 expression (Perincheri et al., 2005). Indeed, closer examination of short-read alignment showed a lack of reads aligning to the start site of the structural variant in BALB/cJ (**Supplementary Figure 5**), thus corroborating the presence of this historical structural variant that is private to BALB/cJ. Raw data can be found at the NCBI Gene Expression Omnibus ascension GSE196352 (striatum) and GSE196334 (hippocampus).

### Proteomic analysis of whole brain tissue from the BALB/c parental substrains corroborates ZHX2 as a functional candidate protein underlying whole brain [OMOR]

Analysis of parental substrain whole brain homogenate protein levels through mass spectrometry revealed 386 differentially expressed proteins with an adjusted p-value < 0.05 and 1377 genes with an unadjusted p-value < 0.05. Of the top 386 proteins (adjusted P < 0.05), only six were contained within the Bayes confidence interval of the chromosome 15 QTL for whole brain [OMOR] (MRPL13, ZHX2, LY6A, BOP1, GCAT, and CYP2D11). Only one of these genes, MRPL13, contains an annotated mutation based on short read sequencing data. Individual expression plots based on Substrain and Sex are shown for ZHX2 (**Figure 4A**), and CYP2D11 (**Figure 4B**). Interestingly, when analyzed separately by sex we observe that female mice show a significant difference and are driving the observed difference (logFC = 0.37, logOdds = 0.97) as opposed to males who did not reach significance (logFC = 0.17, logOdds = -4.37). Complete data set can be found in Beierle et al., 2022.

**Figure 4:**
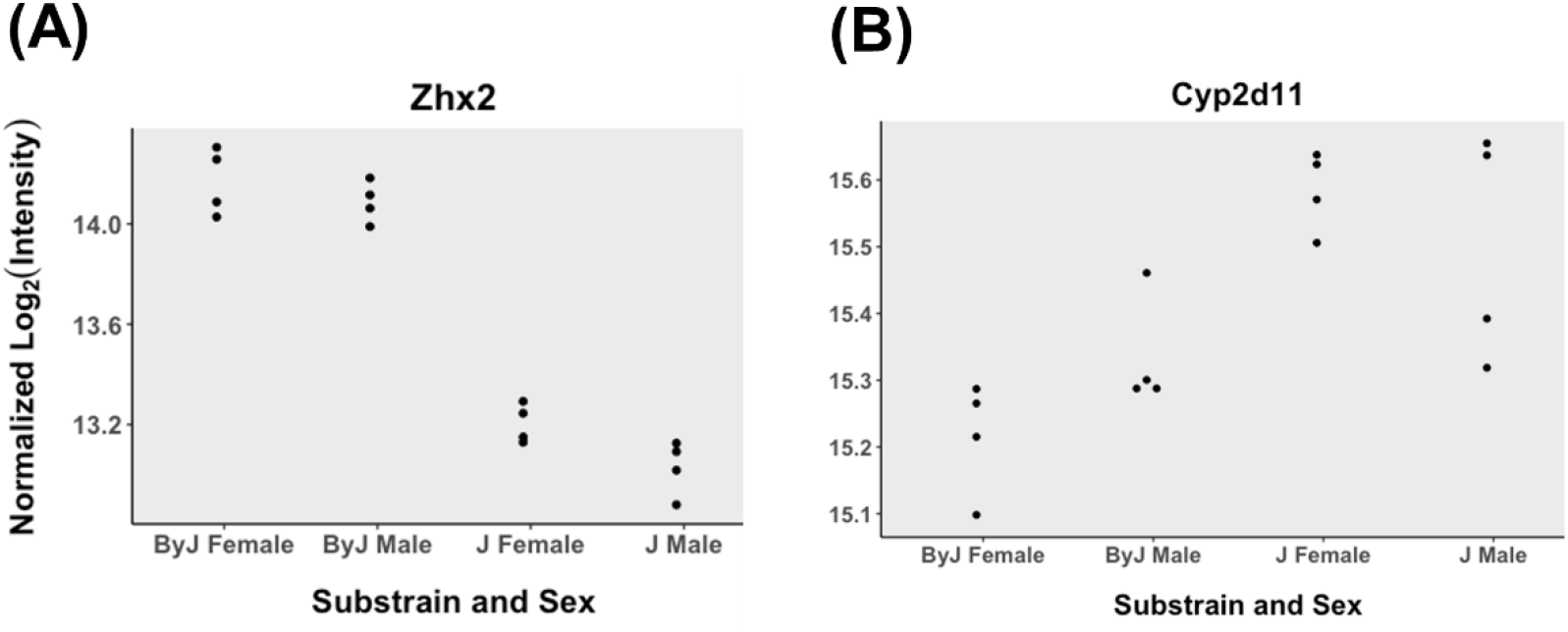
Whole brain proteomics of BALB/c substrains corroborates the Zhx2 eQTL and identifies CYP2D11 as an upregulated protein in J mice. (A): Data generated from 8 BALB/cJ (2/sex/tx) and 8 BALB/cByJ (2/sex/tx) mice. Data from Normalized intensity for Zhx2 in J and By mice (adj pval=2.79e-9). (C): Normalized intensity for CYP2D11 in J and By mice (adj pval=0.0066).

### Transcriptome analysis of liver between BALB/c substrains

Given the identification of a QTL and candidate gene underlying OXY metabolite concentration in the brain, to gain insight into the downstream mechanism, in the absence of having harvested F2 livers, we turned to analysis of the liver in the parental BALB/c substrains which is the primary site of OXY metabolite production. The results of differential gene expression across substrains is shown as a volcano plot in **Figure 5A**. In considering genes with an absolute log2 fold change greater than 0.32 (absolute fold-change > 1.25), we observed a total of 2465 differentially expressed transcripts with an adjusted p < 0.05 (**Supplementary Table 3**). Sixty-one of these transcripts were located within the Bayes QTL interval for brain [OMOR] levels. These genes include Zhx2, but also four Cyp genes, including Cyp2d12, Cyp2d37-ps, Cyp2d40, and CYP2d41-ps which were all downregulated in J mice. Because both the increase in state-dependent OXY-CPP and increase in oxymorphone brain concentration were driven by J females, we also investigated potential sex-specific differential gene expression within the liver (**Supplementary Table 4, 5**). To facilitate legibility, we also plotted female and male-specific alterations in PK related gene expression (defined for this purpose as any CYP, UGT, ABC, or SLC gene and any gene found within the GO terms Xenobiotic Metabolic Process or Xenobiotic transport (GO:0006805, 0042908), **Figure 5B, C**). Sex-stratified analysis identified Ugt2b37 as robustly differentially expressed in females (logFC = - 3.24, adjusted p = 6.96e-05), but not in males (logFC = -0.68, adjusted p = 0.16). Analysis of Ugt2b37 transcript levels revealed a significant main effect of Substrain (F(1, 28)= 20.51, p=1e- 4), Sex (F(1, 28)= 15.82, p=4.4e-4), and an interaction (F(1, 28)=18.07, p=2.1e-4). Tukey’s post hoc test revealed that the interaction was driven by a significant increase in Ugt2b37 in By females versus all other groups (adjP = 1.71e-5 – 6.0e-6), raising the interesting hypothesis that a decrease in Ugt2b37 expression in J females could increase plasma [OMOR] and in turn, increase brain [OMOR]. Raw data can be found at the NCBI Gene Expression Omnibus ascension GSE198375.

**Figure 5:**
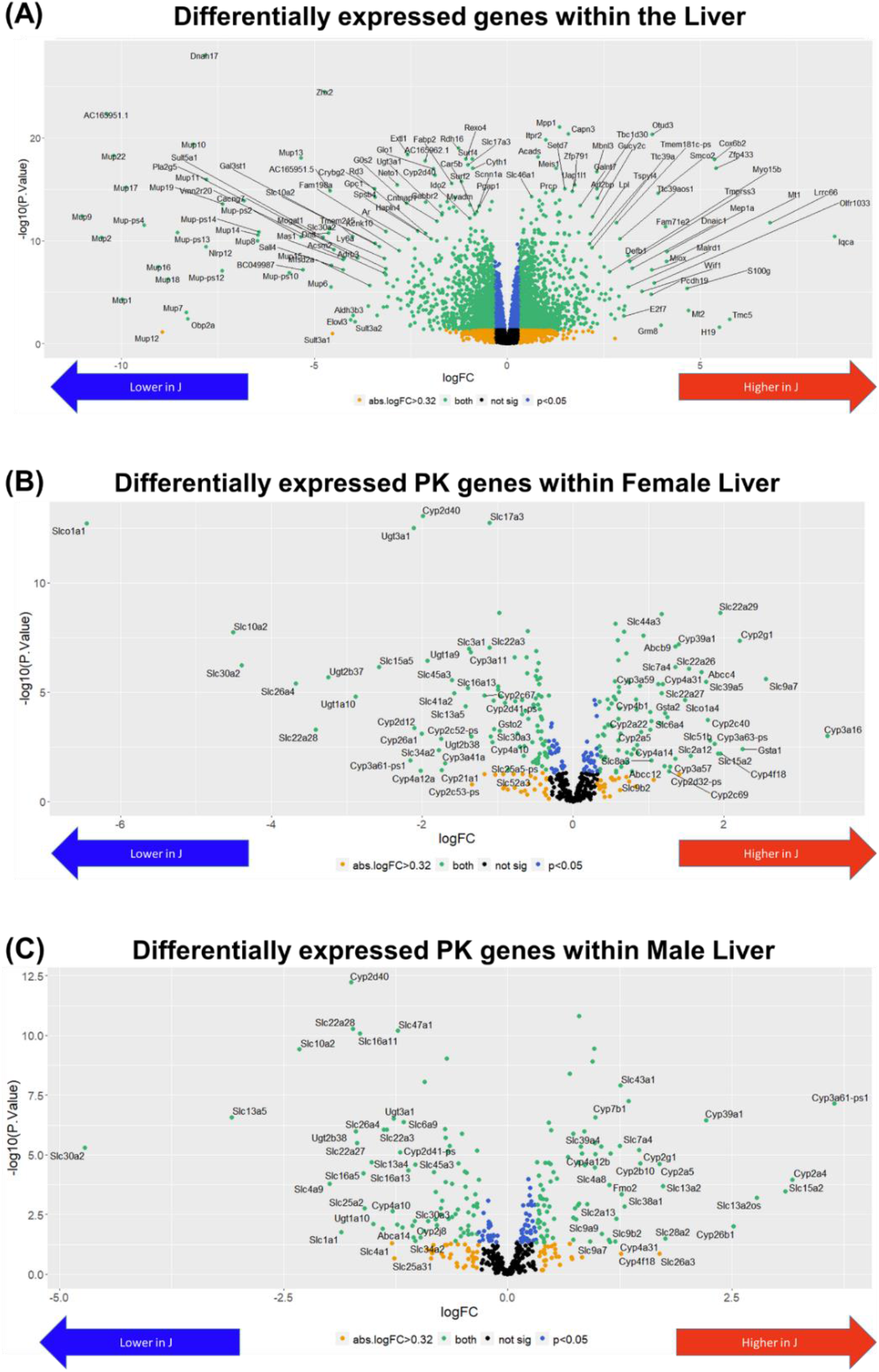
Transcriptome analysis of the liver in BALB/c substrains. **(A):** 16 BALB/cJ (4/sex/tx) and 16 BALB/cByJ mice (4/sex/tx) data were analyzed blocking the effects of sex and treatment. **Sex-combined:** A volcano plot shows the distribution of differentially expressed genes in liver homogenate of all mice (J-By expression). Zhx2 is denoted with an arrow. **(B): Females**: Using only females, data were analyzed blocking the effect of treatment. A volcano plot shows the distribution of differentially expressed PK genes in liver homogenate. For this figure, PK genes include CYP, UGT, ABC, and SLC genes. Note the robust down regulation of Ugtb37 in J females, denoted with an arrow. **(C): Males:** Using only Males, data were analyzed blocking the effect of treatment. A volcano plot shows the distribution of differentially expressed PK genes in liver homogenate of male mice.

### Transcriptome analysis of liver between BALB/c substrains

Analysis of gene expression differences in the liver homogenate protein levels between parental substrains through mass spectrometry revealed 693 differentially expressed proteins with an adjusted p-value < 0.05 and 1667 proteins with an unadjusted p-value < 0.05 (**Figure 6A**). Of the top 693 proteins (adjusted P < 0.05), only 14 were contained within the Bayes confidence interval of the chromosome 15 QTL for whole brain [OMOR] (ZHX2, ZHX1, GM29394, SQLE, NDRG1, ST3GAL1, KHDRBS3, PLEC, LGALS1, PLA2G6, CYP2D11, SERHL, ARFGAP3, and FBLN1). Only one of these genes, KHDRBS3, contains an annotated mutation based on short read sequencing data. Notably, we do observe higher expression of CYP2D11 in J mice (**Figure 6B**). A complete list of differentially expressed liver proteins can be found in **Supplementary Table 6**.

**Figure 6:**
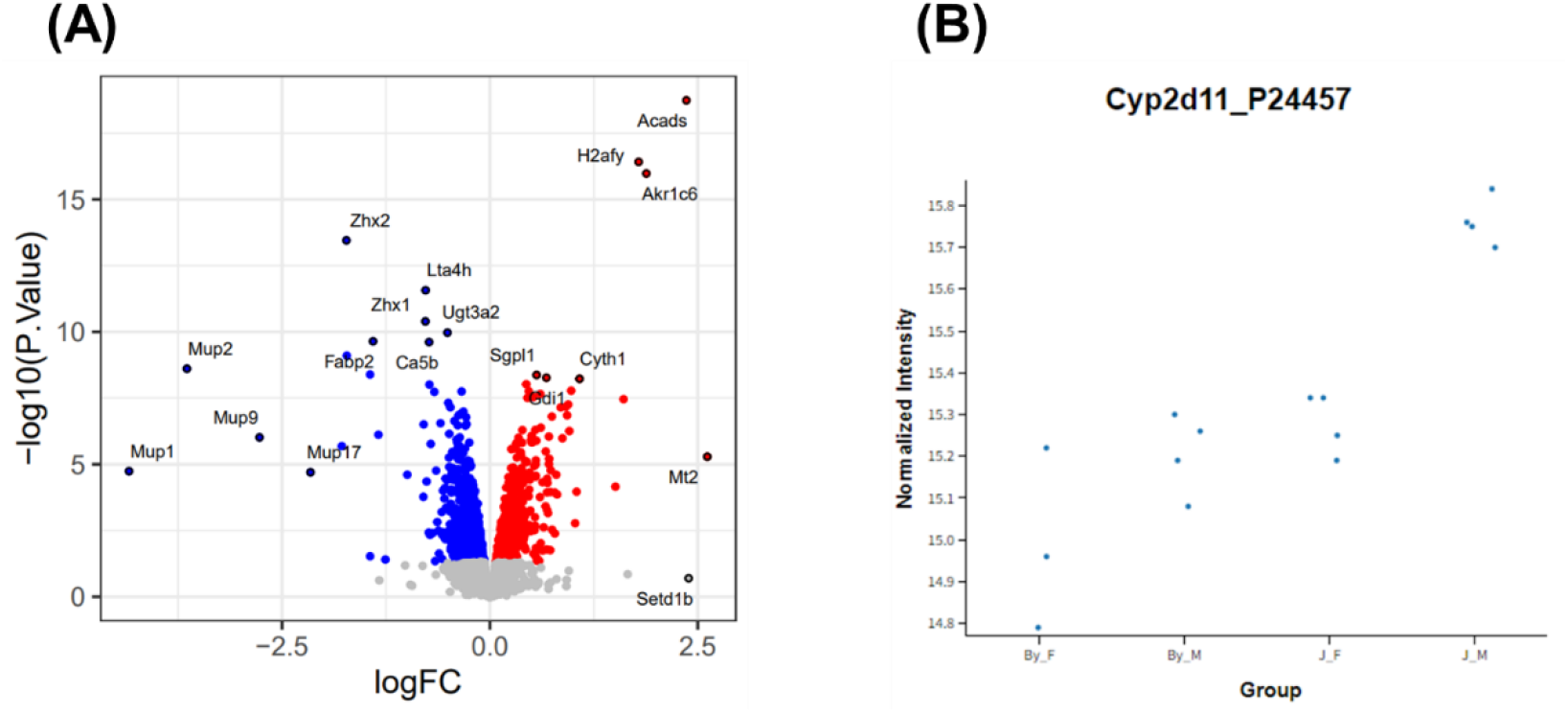
Proteomic analysis of the liver in BALB/c substrains. **(A):** 8 BALB/cJ (4/sex) and 7 BALB/cByJ saline treated mice (3F, 4M) were analyzed. A volcano plot shows the distribution of differentially expressed proteins in liver homogenate of all mice (J-By expression). **(B):** Overlap between differentially expressed RNA transcripts and proteins in the liver. **(C):** Normalized intensity of CYP2D11 across strain and sex.

## Discussion

We identified robust state-dependent CPP in J females and an absence of this behavior in By females, whereas all male mice showed a comparable phenotype (**Figure 1**). A similar trend was observed for OXY-induced locomotion during state-dependent CPP, whereby J females and all male mice showed greater OXY-induced locomotor activity compared to By females (**Figure 1E-G**). Because this substrain difference in behavior was only observed following OXY administration, we wondered whether brain concentration of OXY and/or its metabolites could explain the state-dependent recall of OXY-CPP. We found a robust 1.5 to 2-fold increase in brain [OMOR] in J females at 30 min post-OXY (1.25 mg/kg, i.p.) in two independent experiments (**Figure 2C, F**), demonstrating the robustness of this finding and implicating increased brain [OMOR] as a potential mechanism underlying increased state-dependent OXY-CPP in J females.

While we have not yet made the link between increased brain [OMOR] and increased OXY-induced behavior in J females, it is worthy of further discussion. Because OMOR has a much higher affinity and efficacy at the mu opioid receptor compared to OXY (Peckham & Traynor, 2006; Thompson et al., 2004; Volpe et al., 2011), increased whole brain [OMOR] could facilitate and enhance drug-induced interoception. We hypothesize that increased brain [OMOR] enhances the speed and intensity of the opioid interoceptive effect, that then serves as a more effective unconditioned stimulus, and leads to stronger conditioned stimulus - unconditioned stimulus association with the drug-paired context. A stronger conditioned stimulus - unconditioned stimulus association would sensitize the memory to drug reactivation, increasing state-dependent recall of OXY-CPP.

We identified a single genome-wide QTL on chromosome 15 for whole brain [OMOR (**Figure 3A,B**). Despite parental substrain differences, we did not detect significant QTLs for brain [OXY] or [NOR]. The large genetic effect of the chromosome 15 QTL in (**Figure 3C**) mirrors parental substrain data (**Figure 2D**) and corresponds to a larger portion of the observed female variance compared to the sex-combined analysis (32% vs 19%, **Table 1**). The causal gene/variant could alter production of OMOR (Phase I metabolism by CYP enzymes), degradation of OMOR (Phase II metabolism by UGT enzymes), and/or the transport of OMOR to/from brain (e.g., SLC or ABC transporters).

There is a polymorphic *Cyp2d40* gene and pseudogene *Cyp2d37*-ps within the interval, but both are located 15 Mb from the QTL peak (**Table 2**) and we did not identify any functional eQTL, liver transcriptomic, or proteomic evidence to support them as candidates. Instead, functional eQTL analysis (**Table 2**) combined with proteomics (**Figure 4**) revealed Zhx2 as a candidate gene (**Table 3**). Zhx2 was the only transcript with a cis-eQTL that peaked with the chromosome 15 QTL for brain [OMOR]. Exon-level eQTL analysis revealed the only coding exon (3) within *Zhx2* was differentially used between genotypes at this locus as were three other introns/non-coding exons (**Table 4**). Both whole brain and liver proteomics showed ZHX2 downregulation in J mice, supporting prior studies examining ZHX2 protein in the J substrain (Perincheri et al., 2005). Short read sequencing did not detect variants within or near Zhx2, however, there is a known 6.2 kb mouse endogenous retroviral element (**MERV**) within Zhx2 (Perincheri et al., 2005). Insertion of this MERV leads to the inappropriate splicing of Zhx2 mRNA and decreased protein expression (Perincheri et al., 2008). Our short-read WGS data corroborate the private MERV insertion in the BALB/cJ substrain (**Supplementary Figure 5**). Further details, including MERV DNA sequence, can be found on The Jackson Laboratory Mouse Genome Informatics website.

**Table 2:** Transcript-level cis-eQTLs on chromosome 15 for striatum and hippocampus. Transcripts possessing significant eQTLs on chromosome 15 (FDR < 0.05) in the striatum and hippocampus. Yellow highlighted rows indicate the peak location of each cis-eQTL which in both cases is the same location as the QTL for brain [OMOR]. Analysis was conducted in 64 F2 mice (16/sex/tx), and analysis included Sex, RNA extraction Batch, and Treatment as covariates.

| Region | Gene | Chr | Loc (Mb) | Loc (cM) | pvalue | FDR |
| --- | --- | --- | --- | --- | --- | --- |
| Striatum | Zhx2 | chr15 | 4.31 | 1.95 | 3.1E-07 | 5.7E-03 |
|  | Zhx2 | chr15 | 4.99 | 1.983 | 3.1E-07 | 5.7E-03 |
|  | Zhx2 | chr15 | 7.83 | 3.603 | 2.6E-07 | 4.8E-03 |
|  | Zhx2 | chr15 | 8.12 | 3.702 | 2.6E-07 | 4.8E-03 |
|  | Zhx2 | chr15 | 17.35 | 7.918 | 1.7E-07 | 3.4E-03 |
|  | Zhx2 | chr15 | 21.04 | 8.263 | 6.5E-08 | 1.3E-03 |
|  | Zhx2 | chr15 | 32.32 | 13.546 | 2.5E-13 | 7.3E-09 |
|  | Zhx2 | chr15 | 57.11 | 23.8 | 6.6E-26 | 1.4E-20 |
|  | Zhx2 | chr15 | 81.39 | 38.094 | 1.9E-09 | 4.7E-05 |
|  | Zhx2 | chr15 | 84.44 | 39.864 | 3.9E-08 | 8.0E-04 |
|  | Zhx2 | chr15 | 86.26 | 40.584 | 3.9E-08 | 8.0E-04 |
| Hippocampus | Zhx2 | chr15 | 32.32 | 13.546 | 6.2E-09 | 2.7E-04 |
|  | Zhx2 | chr15 | 57.11 | 23.8 | 1.8E-13 | 7.2E-08 |
|  | Zhx2 | chr15 | 81.39 | 38.094 | 5.4E-09 | 2.4E-04 |
|  | Zhx2 | chr15 | 84.44 | 39.864 | 1.2E-07 | 4.5E-03 |
|  | Zhx2 | chr15 | 86.26 | 40.584 | 1.2E-07 | 4.5E-03 |

**Table 3:** Exon-level cis-eQTLs on chromosome 15 for hippocampus and striatum. Exon-level cis-eQTLs for Zhx2 on (FDR < 0.05) conducted in 64 F2 mice (16/sex/tx), and analysis included Sex, RNA extraction Batch, and Treatment as covariates.

|  | snps | name | Feature | Start (bp) | Length | pvalue | FDR |
| --- | --- | --- | --- | --- | --- | --- | --- |
| Straitum | rs264203947 | Zhx2 | Exon 3 | 57821111 | 2641 | 3.00E-26 | 3.44E-20 |
|  | rs264203947 | Zhx2 | Intron 1 | 57695787 | 66859 | 2.73E-24 | 3.10E-18 |
|  | rs264203947 | Zhx2 | Exon 4 | 57838591 | 1242 | 1.00E-22 | 9.07E-17 |
|  | rs264203947 | Zhx2 | Intron 3 | 57823752 | 14839 | 7.76E-16 | 3.53E-10 |
| H.camp | rs264203947 | Zhx2 | Intron 1 | 57695787 | 66859 | 1.71E-14 | 2.92E-08 |
|  | rs264203947 | Zhx2 | Exon 3 | 57821111 | 2641 | 3.95E-14 | 4.08E-08 |
|  | rs264203947 | Zhx2 | Exon 4 | 57838591 | 1242 | 2.04E-11 | 8.70E-06 |
|  | rs264203947 | Zhx2 | Intron 3 | 57823752 | 14839 | 3.10E-08 | 0.004925 |

Zhx2 (zinc fingers and homeoboxes 2) codes for a globally expressed transcriptional repressor (Kawata et al., 2003). Disruption of Zhx2 expression disrupted expression of hepatic CYP enzymes involved in lipid metabolism, often exhibiting sex-specific changes (Creasy et al., 2016). Zhx2 disruption could also impact the expression of CYPs involved in the metabolism of xenobiotics like oxycodone, given that disruption of central CYP regulators can alter expression of endo- and xeno-biotic metabolizing CYPs (Hu et al., 2011). We hypothesize decreased Zhx2 expression in Js leads to female-specific alterations in the expression of one or more pharmacokinetic genes (e.g., CYP gene(s)) in the brain and/or the liver (e.g., UGT enzymes), culminating in increased brain concentrations of OMOR (**Figure 7**).

**Figure 7:**
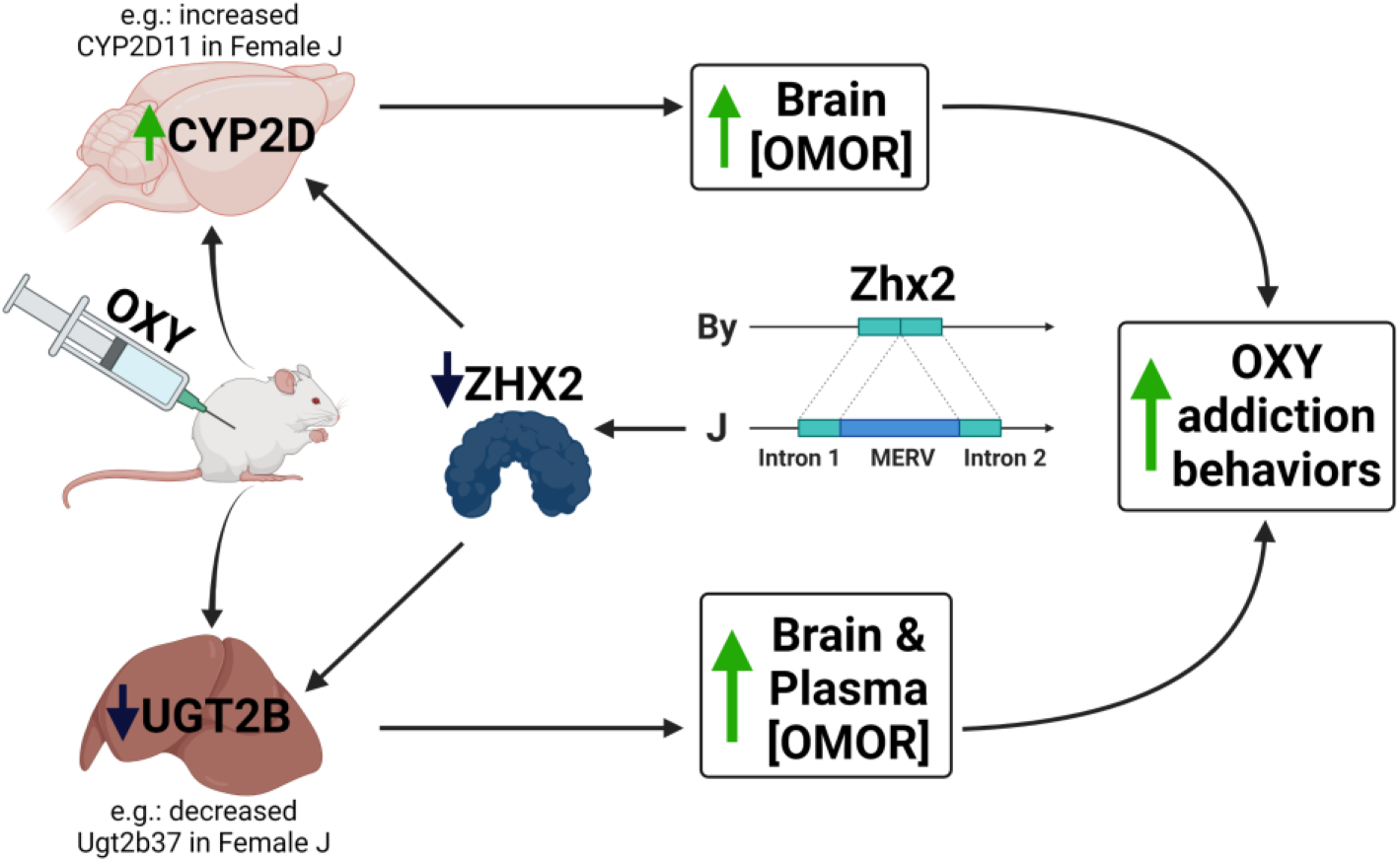
Proposed quantitative trait mechanism linking decreased Zhx2 expression with increased brain concentration of OMOR and increased OXY-induced behaviors. We hypothesize that the insertion of a MERV into Zhx2 leads to decreased protein expression and in turn, changes in expression of liver enzymes that metabolize OXY. The functional consequence is an increase in brain concentration of the highly potent metabolite OMOR which serves as both a potent unconditioned and conditioned stimulus that facilitates both state-dependent learning and state-dependent memory of OXY reward and possibly other OXY-induced behaviors. Created with BioRender.com.

A critical unanswered question is whether the mechanism underlying increased whole brain [OMOR] originates from the liver or the brain. There is a growing appreciation for drug metabolism by CYP enzymes within the brain, including CYP2Ds (Ferguson & Tyndale, 2011; Miksys & Tyndale, 2006; Ravindranath & Strobel, 2013) and manipulation of central CYP2Ds has been shown to modulate OXY, tramadol, and codeine analgesia (McMillan et al., 2019, 2019; McMillan & Tyndale, 2015). Liver transcriptome revealed a pattern of decreased Cyp2d expression in the J substrain, contradicting the hypothesis that increased CYP-mediated OMOR production in the livers of J mice underlies increased brain [OMOR]. However, brain proteomics revealed an upregulation of CYP2D11 protein in Js that was driven by J females, suggesting that increased CYP2D, and increased CYP2D-mediated *brain* production of OMOR could comprise the mechanism. Given Zhx2 is expressed throughout the brain (Ng et al., 2009), Zhx2 it could regulate expression of PK genes within the CNS, including Cyp2d11, leading to increased brain [OMOR]. Notably, liver proteomics revealed CYP2D11 as the only CYP2D gene showing a significant difference in protein and is increased in J mice (**Figure 6B**).

A second hypothesis underlying increased brain [OMOR] is reduced elimination of OMOR, out of the brain or through a direct reduction in UGT-mediated glucuronidation and elimination of OMOR. In support of the latter, we observed a downregulation of *Ugt2b37* RNA in J mice that was much more robust in J females compared to J males (Female logFC = -3.24, Male logFC = -0.68). *Ugt2b37* is a UGT enzyme that mediates phase 2 glucuronidation of morphine in mice (Kurita et al., 2017). Reduced OMOR conjugation would decrease OMOR clearance, increase plasma OMOR, and increase brain [OMOR]. However, differences in expression were driven by By female mice, and did not mirror the patterns observed in brain [OMOR], where J females comprised the differences. Additionally, we could not confirm these differences at the proteomics level, as we were unable to detect UGT2B37 within the liver.

There are limitations to this study. First, our metabolite profiling relies on a single dose, time point, route of administration, and tissue. The reason stems from the original intent of this study: to map the genetic basis of state-dependent OXY-CPP. Full tissue-concentration curves (e.g., brain versus plasma) are necessary to inform the underlying pharmacokinetic mechanism. Second, we were unable to identify QTLs underlying state-dependent learning of OXY reward. The heritability of CPP is notoriously low (alcohol, Cunningham et al., 1991; methamphetamine in LG and SM strains, Bryant et al., 2014; cocaine in BXD Philip et al., 2010) due mainly to the large environmental variance, necessitating a much larger sample size for detecting a QTL. Based on parental strain effect sizes, we would have needed to test 311 OXY treated F2 mice to detect QTLs with 80% power (622 with saline controls). Rather than test more F2 mice, we will test the effect of Zhx2 on OXY behavior in future gene editing studies focused on the quantitative trait mechanism linking Zhx2 to brain [OMOR].

This study highlights the importance of assessing pharmacokinetic measures in genetic analysis of drug-induced behaviors. It also highlights how Sex can dramatically alter brain metabolite concentration in a genotype-dependent manner, which is important given sex differences in substance use disorders (Cornish & Prasad, 2021) including opioid use disorder (Chartoff & McHugh, 2016). This study also highlights the limitations of relying solely on short read DNA sequencing to annotate genetic variants and the utility of including proteomics in genetic regulation of gene expression associated with complex traits. Without prior literature, we would have missed a critical functional variant underlying differences in Zhx2 expression. Finally, this study highlights the power of reduced complexity crosses for rapidly identifying functional candidate genes underlying complex traits (Bryant et al., 2018; 2020).

In summary, we identified a QTL containing a highly plausible candidate gene underlying whole brain [OMOR]. We hypothesize that increased brain [OMOR] brain in females is mediated by a MERV-mediated loss of Zhx2 function (**Figure 7**). Furthermore, we hypothesize that dysregulation of one or more proteins involved in OXY pharmacokinetics increases whole brain [OMOR] and state-dependent OXY reward learning (**Figure 7**). Future studies will test these hypotheses using reciprocal gene editing on the two BALB/c genetic backgrounds and generating full tissue-concentration profiles of OXY and metabolites in these lines. We will also use tissue-specific viral targeting to identify relevant tissue(s) underlying increased brain [OMOR] concentration. Furthermore, it will be important to test the role of Zhx2 in the pharmacokinetic profiles of other opioid drugs, addiction-relevant model behaviors, and antinociception. Finally, given the nominal association of the ZHX2 locus with the quantity and frequency of nicotine use in humans (p = 7 x 10^-7^) (McGue et al., 2013), we will test the influence of Zhx2 in the pharmacokinetic profile and addiction liability to other drugs of abuse.

## Acknowledgments

Analytical metabolite mass spectrometry work was performed by the Center for Human Toxicology through NIDA Project N01DA-19-8951 (contract# 75N95019C00016).

This research was supported by generous ongoing support from BU to AE and the CNSB.

## Footnotes

This research was funded by the National Institute of Health’s National Institute on Drug Abuse [U01DA050243, R01DA039168, 5U01DA04439902]; The national Institute of Allergy and Infectious disease [U19 AI100625, P01 AI132130] The National Institute of General Medical Sciences [T32GM008541]; and the Burroughs Welcome Fund Transformative Training Program in Addiction Science [1011479].

## Authorship contributions

*Participated in study design*: Beierle JA, Bryant CD, Ferris MT

*Provided whole genome sequence of BALB/cByJ mice*: Ferris MT

*Conducted Parental strain experiments*: Beierle JA, Yao EJ, Scotellaro JL

*Conducted F2 experiments*: Beierle JA

*Analyzed behavioral video recordings*: Beierle JA, Yao EJ, Scotellaro JL, Sena KD, Wong AL

*Conducted additional genotyping*: Beierle JA, Sena KD

*Conducted RNA extraction, RNA-seq analysis, QTL, eQTL, and other statistics*: Beierle JA

*Conducted Oxycodone and metabolite Mass Spectrometry and analysis*: Averin O, Moody DE, Reilly CA

*Conducted Proteomics Mass Spectrometry and analysis*: Goldstein SI, Lynch WD, Emili A

*Wrote or contributed to the manuscript*: Beierle JA, Bryant CD, Reilly CA, Peltz G, Emili A, Ferris MT

