## Supplemental Figures and Legends for "Zhx2 is a candidate gene underlying oxymorphone metabolite brain concentration associated with state-dependent oxycodone reward"

### Supplementary Figure Legends

#### Supplementary Figure 1: Drug-Free OXY-CPP in BALB/c substrains

**(A)**: Results were analyzed using a three way ANOVA considering Substrain, Treatment, and Sex, error bars represent standard deviation. CPP was assessed in the drug-free state on Day 8 following an injection of SAL (i.p.), there was an effect of Treatment ($F(1, 276)= 5.58$, $p=0.019$), but no effect of Substrain ($F(1, 276)= 1.9$, $p=0.17$), Sex ($F(1, 276)= 1.91$, $p=0.17$), or any interactions [Treatment x Substrain ($F(1, 276)= 0.11$, $p=0.74$); Treatment x Sex ($F(1, 276)= 0.002$, $p=0.96$); Substrain x Sex ($F(1, 276)= 2.43$, $p=0.12$); Treatment x Substrain x Sex ($F(1, 276)= 0.35$, $p=0.56$)].

#### Supplementary Figure 2: Breeding, genotyping, and phenotyping of F2 mice.

**(A):** Genetic map of the 218 miniMUGA markers and 1 added marker used for QTL mapping. **(B):** Three way ANOVA considering Sex, Cohort, and Age of F2 mice indicated a main effect of Sex on whole brain [OMOR] ($p=5.4e-5$). Thus Sex was included as an additive covariate in the QTL model.

#### Supplementary Figure 3: Day 9 QTL mapping measures

**(A):** QTL plot demonstrating no significant linkage between genotype and Day 9 phenotypes when considering treatment as an interactive covariate and sex as an additive covariate. N= 200 OXY, 177 SAL. **(B):** Effect plot at trending chr10 QTL marker of D9 preference showed an association driven by heterozygous animals. **(C):** Female-specific QTL plot demonstrating no significant linkage between genotype and Day 9 phenotypes when considering treatment as an interactive covariate. N= 108 OXY, 93 SAL. **(D):** Male specific QTL plot demonstrating no significant linkage between genotype and Day 9 phenotypes when considering treatment as an interactive covariate. N=92 OXY, 84 SAL.

**Supplementary Figure 4: Power analysis for QTL analysis of brain OMOR concentration and state-dependent OXY-CPP. (A):** Power analysis showed that, with our sample size (N=133), we were powered to detect QTLs explaining 14% of the phenotypic variance or more. **(B)**: Power analysis showing, with the observed state dependent CPP effect size and variation in parental strain mice, we need to test 311 OXY treated F2 mice to be 80% powered to detect QTLs. **(C)** Given our 200 OXY treated mice tested for D9, we are powered to detect QTLs explaining ~12% of the observed variance in our F2 cohort.

**Supplementary Figure 5: Whole genome sequencing short read alignment at Zhx2 MERV insertion site.** A lack of read alignment at the start site for the MERV insertion was observed in the BALB/cJ substrain known to possess the MERV. In contrast, read alignment was observed in the other two substrains, namely BALB/cJBomTac and BALB/cByJ substrains.

### **Supplementary Tables:**

| | |
|---|---|
| SNP information | >mm10_snp142_rs264203947 range=chr15:57105089-57105129 5'pad=20 3'pad=20 strand=+ repeatMasking=none<br><br>TAGAACATTACTGTTAGTCT**G**ATAACAGTAAAAGAGATGAC |
| SNP sequence +/- 300 bp: Used for genotype | >mm10_snp142_rs264203947 range=chr15:57104809-57105409 5'pad=300 3'pad=300 strand=+ repeatMasking=none<br>TTGGCTGCATGCCCTCTTTCACCATAGTCTTGGTATGTGTGCGTGCATGT<br>GTGCGTATGAGTGTTCATACAGAGCCCACTGGTGATGTAATACTCGCCTT<br>CCTTTGTGTCTCAACTGACTGAAAAGTGGCCAGATGAATAATTTGGCACC<br>ATTTCTTTGTGGTTTTCTCTGAAGCTTGCTACCAGAGAGTCATCTAATTC<br>TTCTTCACTCTGAATCAAGCATTGTAGGCAGGAATATTCTTTAGTTCAAT<br>ATGGTATTCAGTCAACTGAAAACACTCTTATAGAACATTACTGTTAGTCT<br>**G**ATAACAGTAAAAGAGATGACATTGGTAGACATAGAAATTGGAATGATTT<br>ATAAATGTACACTTCGTGCTCCTTTGAAGTGGATTTCCATGCAAGAGGGA<br>AGAATATCAAAGCCCTTGAAGTTCAGTGCTGCTCCTTCAAGCTCATGAAT<br>TAAGAACTTTTAGAGTTGCCCAGAGCTTTGTTATTATTATAACAATATAT<br>TATTATTATTTCTGCTTTCATGTGCTATCATCTCCTGTTTTAACAAATCT<br>TTAAGGAAGAATTTCTCATTTCCTACCTTTGTCCCATTTTGAAGGTAGTG<br>A |
| SNP sequence +/- 300 bp: Used for | TTGGCTGCATGCCCTCTTTCACCATAGTCTTGGTATGTGTGCGTGCATGTG<br>TGCGTATGAGTGTTCATACAGAGCCCACTGGTGATGTAATACTCGCCTTCC<br>TTTGTGTCTCAACTGACTGAAAAGTGGCCAGATGAATAATTTGGCACCATT<br>TCTTTGTGGTTTTCTCTGAAGCTTGCTACCAGAGAGTCATCTAATTCTTCT<br>TCACTCTGAATCAAGCATTGTAGGCAGGAATATTCTTTAGTTCAATATGGT<br>ATTCAGTCAACTGAAAACACTCTTATAGAACATTACTGTTAGTCT**[G/T]**A<br>TAACAGTAAAAGAGATGACATTGGTAGACATAGAAATTGGAATGATTTATA<br>AATGTACACTTCGTGCTCCTTTGAAGTGGATTTCCATGCAAGAGGGAAGAA<br>TATCAAAGCCCTTGAAGTTCAGTGCTGCTCCTTCAAGCTCATGAATTAAGA<br>ACTTTTAGAGTTGCCCAGAGCTTTGTTATTATTATAACAATATATTATTAT<br>TATTTCTGCTTTCATGTGCTATCATCTCCTGTTTTAACAAATCTTTAAGGA<br>AGAATTTCTCATTTCCTACCTTTGTCCCATTTTGAAGGTAGTGA |

**Supplementary Table 1: Genotyping marker information**

Sequence used to design Taqman fluorescent genotyping probe. Polymorphic nucleotide is indicated by underline.

**Supplementary Table 2: Polymorphic genes within chr15 QTL**

List of genetic polymorphisms within the chromosome 15 QTL region.

**Supplementary Table 3: Liver RNA-seq DEG list**

List of differentially expressed genes at the RNA level in parental strain livers (BALB/cJ as ref.).

**Supplementary Table 4: Female Liver RNA-seq DEG list**

List of differentially expressed genes at the RNA level in female parental strain livers BALB/cJ as ref.).

**Supplementary Table 5: Male Liver RNA-seq DEG list**

List of differentially expressed genes at the RNA level in male parental strain livers BALB/cJ as ref.).

**Supplementary Table 6: Liver proteomic DEG list**

List of differentially expressed genes at the protein level in parental strain livers BALB/cJ as ref.).

**Supplementary Figures:**

**Supplementary Figure 1: Drug-Free OXY-CPP in BALB/c substrains**

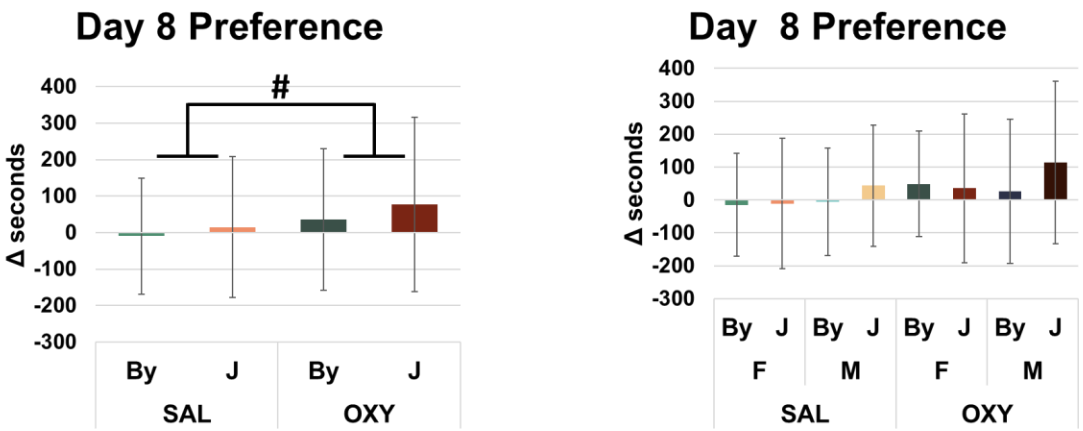


**Supplementary Figure 2: Breeding, genotyping, and phenotyping of F2 mice.**

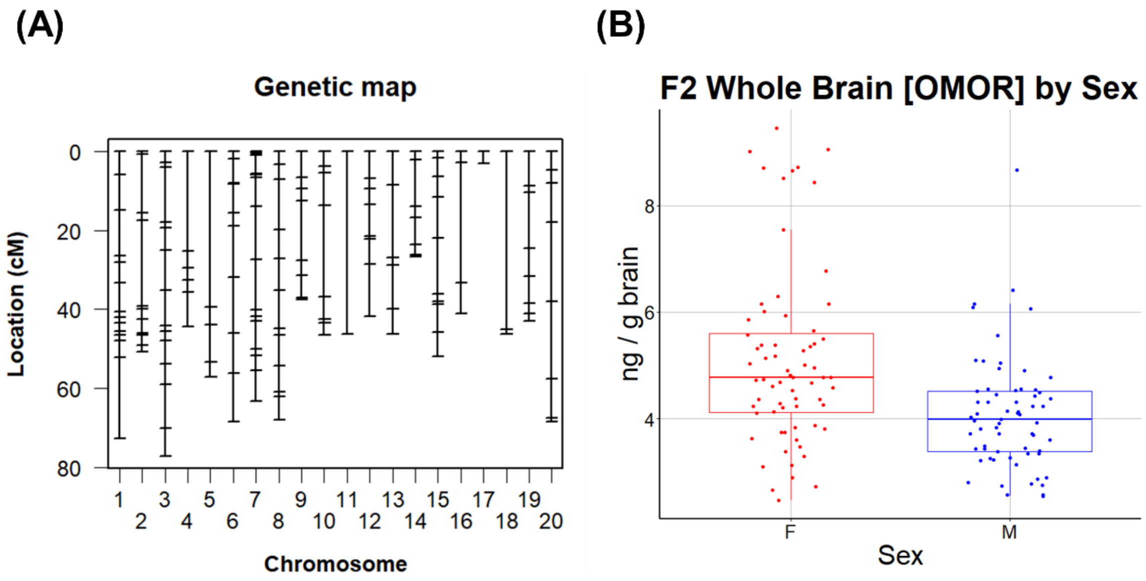

### Supplementary Figure 3: Day 9 QTL Mapping Measures

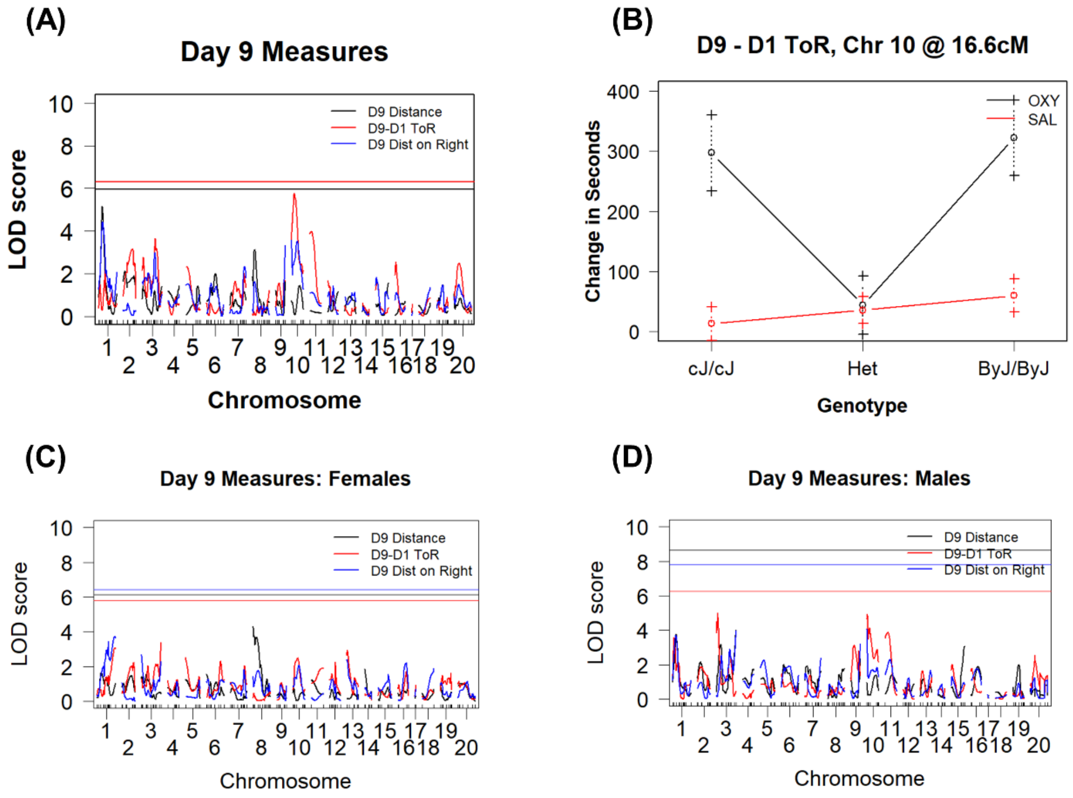


### Supplementary Figure 4: Power analysis for QTL analysis of brain [OMOR] and state-dependent OXY CPP

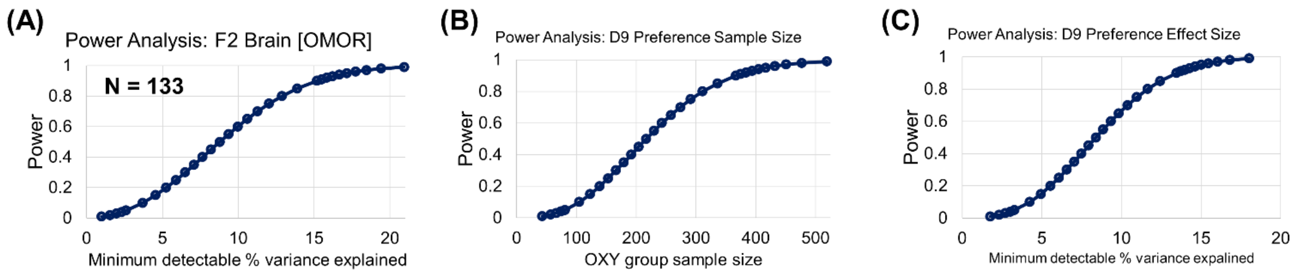

**Supplementary Figure 5: Whole genome sequencing short read alignment at Zhx2 MERV insertion site**

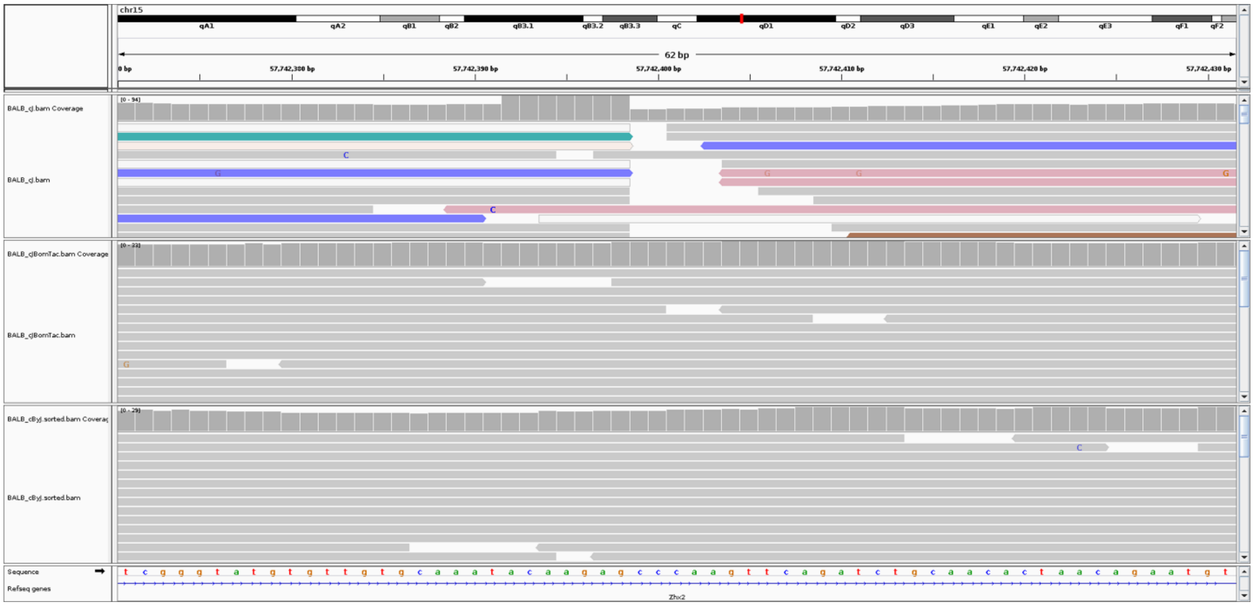
